# Improving Cell-type-specific 3D Genome Architectures Prediction Leveraging Graph Neural Networks

**DOI:** 10.1101/2024.05.21.595047

**Authors:** Ruoyun Wang, Weicheng Ma, Aryan Soltani Mohammadi, Saba Shahsavari, Soroush Vosoughi, Xiaofeng Wang

## Abstract

The mammalian genome organizes into complex three-dimensional structures, where interactions among chromatin regulatory elements play a pivotal role in mediating biological functions, highlighting the significance of genomic region interactions in biological research. Traditional biological sequencing techniques like HiC and MicroC, commonly employed to estimate these interactions, are resource-intensive and time-consuming, especially given the vast array of cell lines and tissues involved. With the advent of advanced machine learning (ML) methodologies, there has been a push towards developing ML models to predict genomic interactions. However, while these models excel in predicting interactions for cell lines similar to their training data, they often fail to generalize across distantly related cell lines or accurately predict interactions specific to certain cell lines. Identifying the potential oversight of excluding example genomic region interaction information from model inputs as a fundamental limitation, this paper introduces GRACHIP, a model rooted in graph neural network technology aiming to address this issue by incorporating detailed interaction information as a hint. Through extensive testing across various cell lines, GRACHIP not only demonstrates exceptional accuracy in predicting chromatin interaction intensity but showcases remarkable generalizability to cell lines not encountered during training. Consequently, GRACHIP emerges as a potent research tool, offering a viable alternative to conventional sequencing methods for analyzing the interactions and three-dimensional organization of mammalian genomes, thus alleviating the dependency on expensive and time-consuming biological sequencing techniques. It also offers an alternative way for researchers to investigate 3D chromatin interactions and simulate their changes in model systems to test their hypotheses.

## Introduction

In mammalian genomes, three-dimensional architectures of chromatin play an important role in regulating gene expression, cellular differentiation, and multiple biological processes[1]. Specifically, chromatin folds into topological associating domains (TADs)[2] and restricts enhancer-promoter interactions. The genome architectures can be captured through genome-wide sequencing assays such as Hi-C, micro-C, HiChIP, etc[3–5]. However, The genome organization is cell-type specific, and high-resolution sequencing of the 3D genome for numerous cell lines is both costly and time-consuming. In addition, high sequencing depth is usually required to capture more interactions, making this process even more costly.

With the prosperity of development in the field of machine learning (ML), research has been conducted to predict genomic interactions leveraging ML models. Most such models build their predictions upon DNA sequence information and genomic features, e.g., Akita[6] and Orca[7]. The use of DNA sequence information is rooted in the fact that DNA sequences contain key protein-binding motifs influencing chromatin folding and overall genome structure. However, a significant challenge with relying chromatin interaction predictions solely on DNA sequence data is the difficulty in generalizing models across different tissue types, given the cell type-specific nature of chromatin interactions, whereas DNA sequences are largely consistent. To overcome this, genomic features are frequently incorporated into predictive models. Commonly used features include ATAC-seq/DNase-seq [8, 9] and CTCF ChIP-seq signals [8], which help identify open chromatin and binding sites of the CTCF protein. Additionally, histone modifications such as H3K4me and H3K27ac are utilized to reflect active regulatory regions, contributing to the cell type-specific information critical for understanding genome organization. Moreover, the incorporation of short-distance co-occurrence information—where genomic regions with distinct genomic features and DNA sequences are closely situated—has proven to be valuable in predicting 3D genome structures. This has led advanced models, such as c.origami [8], to employ convolutional neural networks (CNNs) [10] to unravel these intricate patterns.

Despite the efficacy of these models within the same or analogous cellular environments as their training datasets, their applicability to genetically or functionally diverse cell lines remains constrained. For instance, significant disparities in performance are evident when comparing the efficacy of c.origami, a leading model in chromatin interaction prediction, across the GM12878 cell line and either the H1 or HFF cell lines. In addition, c.origami tends to give false positive predictions at specific genomic regions.

The Results section illustrates both observations in full detail. This limitation in generalizability may stem from an inherent oversight in model design, specifically neglecting the influence of known chromatin interaction intensity on the prediction of genomic interactions. More explicitly, the propensity of certain genomic regions to associate with others distinguishes them from counterparts that, despite sharing similar DNA sequences and multi-omic characteristics, do not exhibit the same interactive patterns—a critical aspect unaddressed by existing models. Justifying our consideration, prior research has established that interactions both between pairs of promoters or enhancers and across multiple cis-regulatory regions (often described as enhancer hubs or enhancer communities) [11, 12] are essential for transcription activation.

To mitigate the limitations of existing models and enhance the precision of genomic interaction forecasts, this study introduces an approach that incorporates chromatin interaction intensity data into the ML model. The model proposed in this paper is augmented with genomic interactions derived from a specific cell line H1 to bolster performance across a variety of cell lines. To achieve this aim, the model is fed genomic interaction data formatted as a sparse graph. A graph neural network (GNN) is then employed to encode this genomic interaction information, thereby refining the representations of genomic regions based on their connections within the graph. Complementarily, the model integrates Transformer [13] and CNN encoding blocks to further refine and condense information from genomic region representations, aiming at elevating the accuracy of predictions. DNABert [14] is applied to encode DNA sequence information, enabling the introduction of more nuanced information derived from DNA sequences and achieving swifter DNA sequence processing compared to models employing one-hot DNA encodings, such as c.origami. This model, termed the Graph-based Chromatin Interaction Prediction model (GRACHIP), is elaborately described in the Model Structure subsection under the Methods section.

GRACHIP has established itself as a robust predictive model for chromatin interactions, validated through comprehensive evaluations across a diverse range of cell lines. This model leverages essential insights together with a pioneering pre-training method to deliver outstanding zero-shot generalizability to previously unobserved cell lines. Remarkably, GRACHIP achieves this using only ATAC-seq and a minimal set of epigenomic signals, thereby streamlining the data requirements and significantly enhancing the feasibility of conducting biological research on chromatin interactions. Furthermore, GRACHIP provides a cost-effective platform for formulating new hypotheses about chromatin interactions, effectively bypassing the substantial expenses typically involved in chromatin conformation capture experiments. This makes GRACHIP an invaluable tool in the field of genomic research, enabling studies that were previously impractical due to cost or resource limitations.

## Results

### GRACHIP accurately predicts genomic interactions

This paper presents GRACHIP, a deep-learning model for predicting the 3D genomic interactions based on DNA sequence information and genomic features, hinted by chromatin interaction intensity information. GRACHIP utilizes DNABert for encoding DNA sequences into more compact, lower-dimensional formats, thereby facilitating more efficient computation while preserving the essential co-occurrence patterns found within and between DNA motifs. The genomic features used by GRACHIP include ATAC-seq and five CUT-RUN features (i.e., CTCF, H3K4me3, H3K4me, H3K27ac, and H3K27me). Fig. 1d illustrates the full process of how the input to GRACHIP is shaped. To assimilate chromatin interaction intensity data into the model, a subset (20%) of this information from the H1 HiC dataset is randomly selected and formatted into *n × n* matrices where *n* denotes the number of genomic regions encompassed per instance. These matrices are then incorporated into the GCN encoder module as edge attributes. Positional encoding akin to BERT [15], a Transformer-based text processing model, is added to the representations of genomic regions within each instance to further enhance the position and order information among genomic regions in each input window. GRACHIP is configured to predict predictions in 2Mb windows with a bin size of 10kb to cover most intra-chromosome interactions. The model structure and the functionality of the underlying network architectures are detailed in the Methods section.

**Fig. 1:**
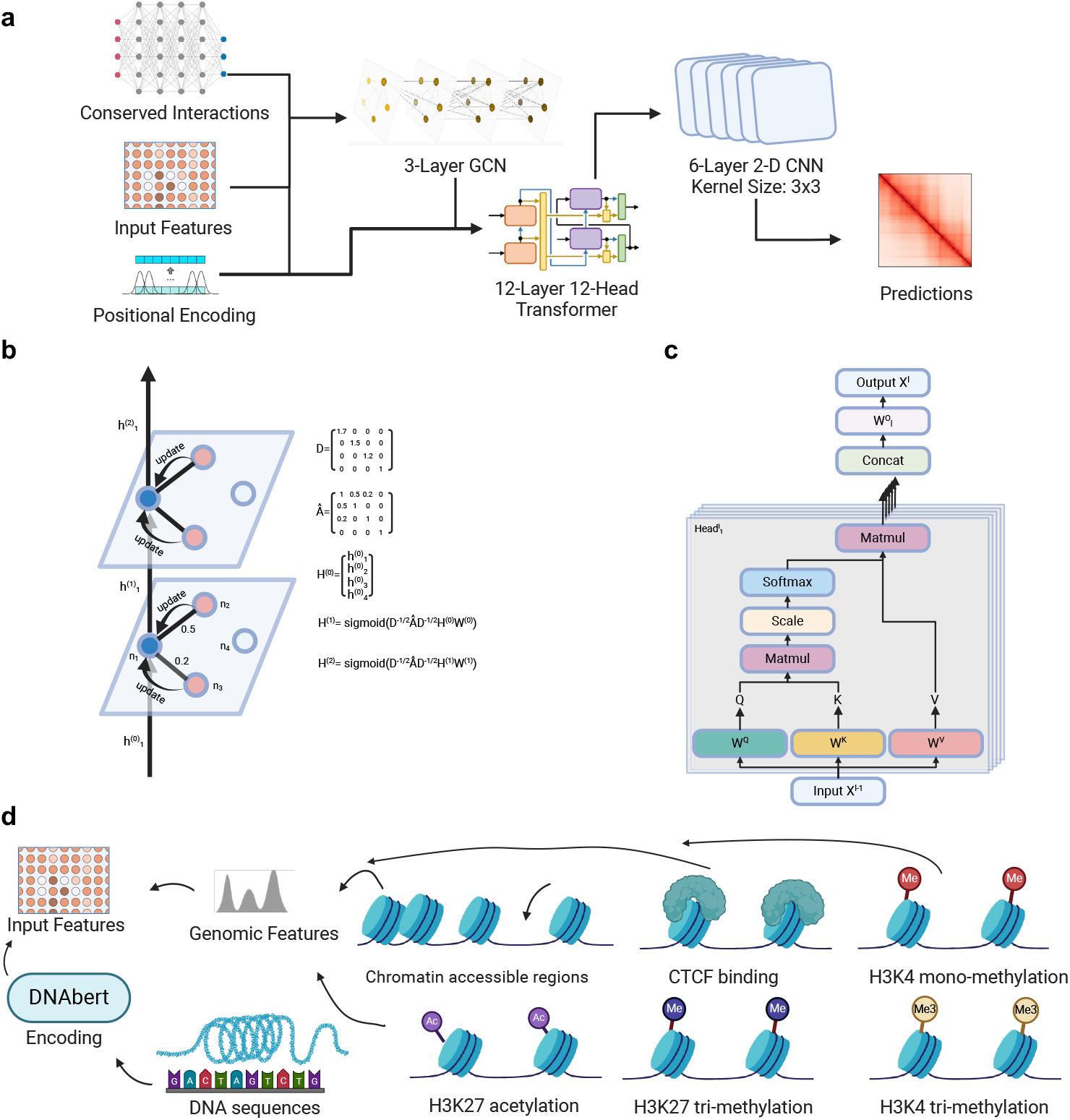
Overview of GRACHIP’s Design and Input Processing. a, the architectural framework of the proposed GRACHIP model. b, an illustration of the GCN mechanism for updating genomic region encodings with chromatin interaction intensity data. c, a demonstration of the Transformer encoder block utilized within GRACHIP. d, an illustration of the composition of GRACHIP’s input, encompassing DNA sequences processed by DNABert alongside various genomic features.

The pre-training of GRACHIP is performed on 20 chromosomes from the HiC-generated interaction matrix of the H1 and HFF cell lines, while Chr10 and chr15 are left out for evaluation purposes only.

In these experiments, GRACHIP consistently delivers exceptional performance, achieving metrics that exceed 90% across Pearson and Spearman correlation coefficients, as well as insulation index correlation scores. This highlights GRACHIP’s robustness in accurately predicting chromatin interactions. To demonstrate the model’s precision, we present two exemplary cases from the H1 and HFF cell lines (Fig. 2a-f). Despite significant discrepancies between the actual labels and the genomic interaction intensity data (derived solely from the H1 HiC dataset), GRACHIP adeptly recovers the target labels in these examples.

**Fig. 2:**
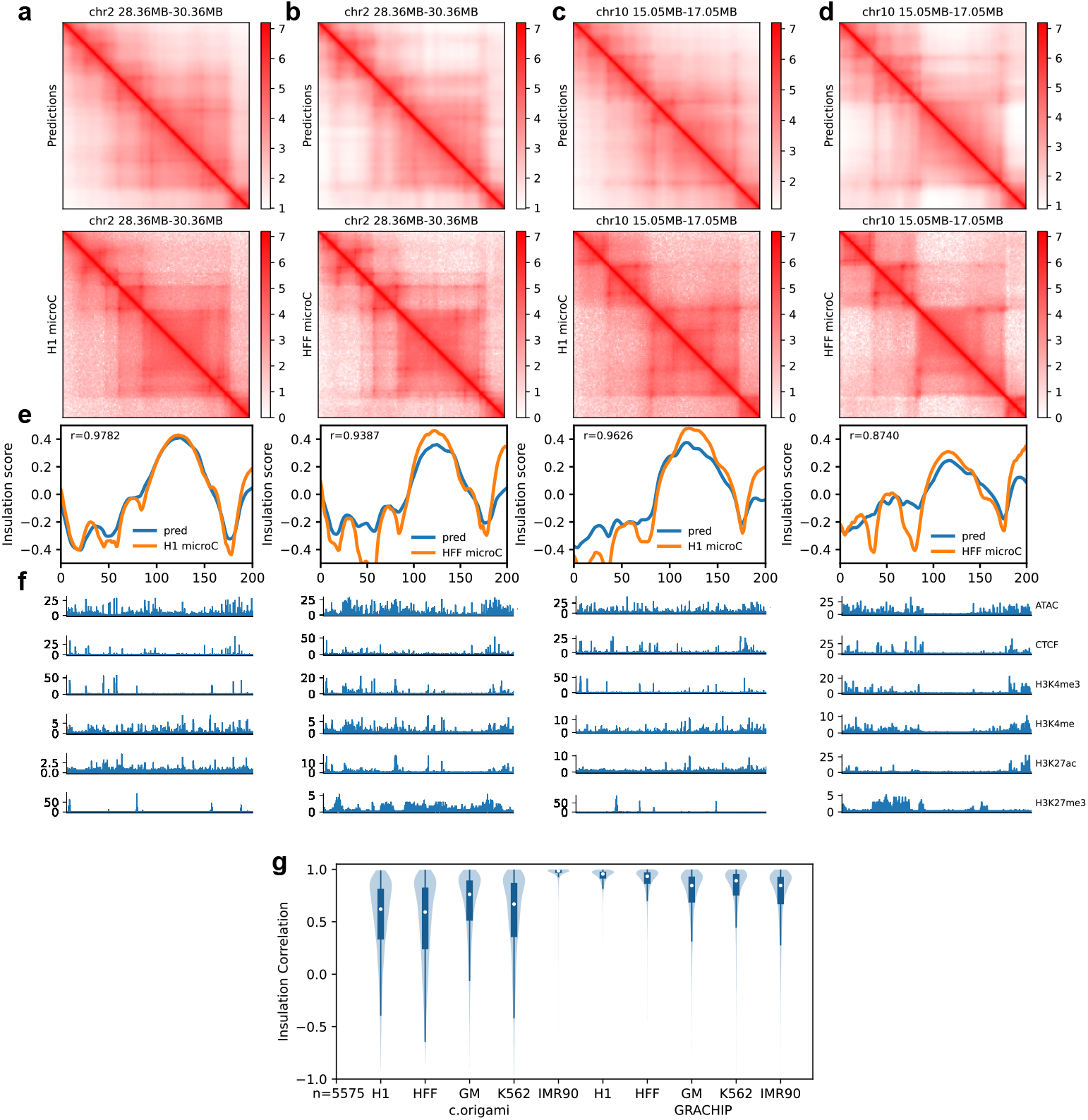
GRACHIP accurately recovers genomic interaction maps. a-d, Predicted genomic interaction (top), true genomic interaction (middle) on H1 and HFF cell line. e, insulation index of the H1, HFF and predicted interaction matrices. f, Genomic features of shown genomic regions. g, Genome-wide correlation of insulation scores across 5 cell types, i.e., H1, HFF, GM12878, K562, and IMR90. The left group of four violins represents predictions made by c.origami, while the right group of four violins shows predictions by GRACHIP.

Remarkably, GRACHIP maintains high performance across all evaluated cell lines, including H1, HFF, GM12878, K562, and IMR90. Fig. 2g visualizes GRACHIP’s superior performance across the whole genome compared to c.origami, where GRACHIP shows considerable advantages. The performance differential is notably significant in GM12878 and K562, two cell lines different from the cell lines used in the training of GRACHIP and c.origami, with GRACHIP achieving median insulation correlations that are 0.08 and 0.13 higher respectively than those of c.origami, transitioning from moderate to strong correlations [16].

While c.origami performs exceptionally well on its training cell line (IMR90) with a median insulation correlation of 0.99, its performance on GRACHIP’s training cell lines (H1 and HFF) is much lower, with median insulation correlations of 0.62 and 0.59, respectively, categorized as moderate correlations. In contrast, GRACHIP achieves median insulation correlations ranging from 0.84 to 0.96 across all five cell lines, indicating strong to very strong correlations. This underscores GRACHIP’s substantial capability and high generalizability across diverse cell lines. Detailed cell type-specific assessments are further explored in the following section.

### Cell-type-specific genomic contact map prediction

In addition to assessing the overall performance of GRACHIP across various datasets, we conducted detailed analyses on genomic regions that show deviations from the training data. We compared the predictions of GRACHIP and c.origami in a representative region on chromosome 2 across four cell lines, as illustrated in Fig. 3a-l. Here, GRACHIP consistently outperforms c.origami, achieving high prediction accuracy. Notably, GRACHIP maintains insulation correlations exceeding 0.85, as evidenced by Pearson correlation coefficients (see Fig. 3m and Fig. S1). This level of consistency is also observed in regions that significantly diverge from the model’s training data (see Fig. 3a-d), highlighting GRACHIP’s remarkable generalizability. Consequently, GRACHIP serves as an invaluable tool for researchers exploring chromatin interactions in less commonly studied cell lines, providing robust predictions without the need for re-training or fine-tuning.

**Fig. 3:**
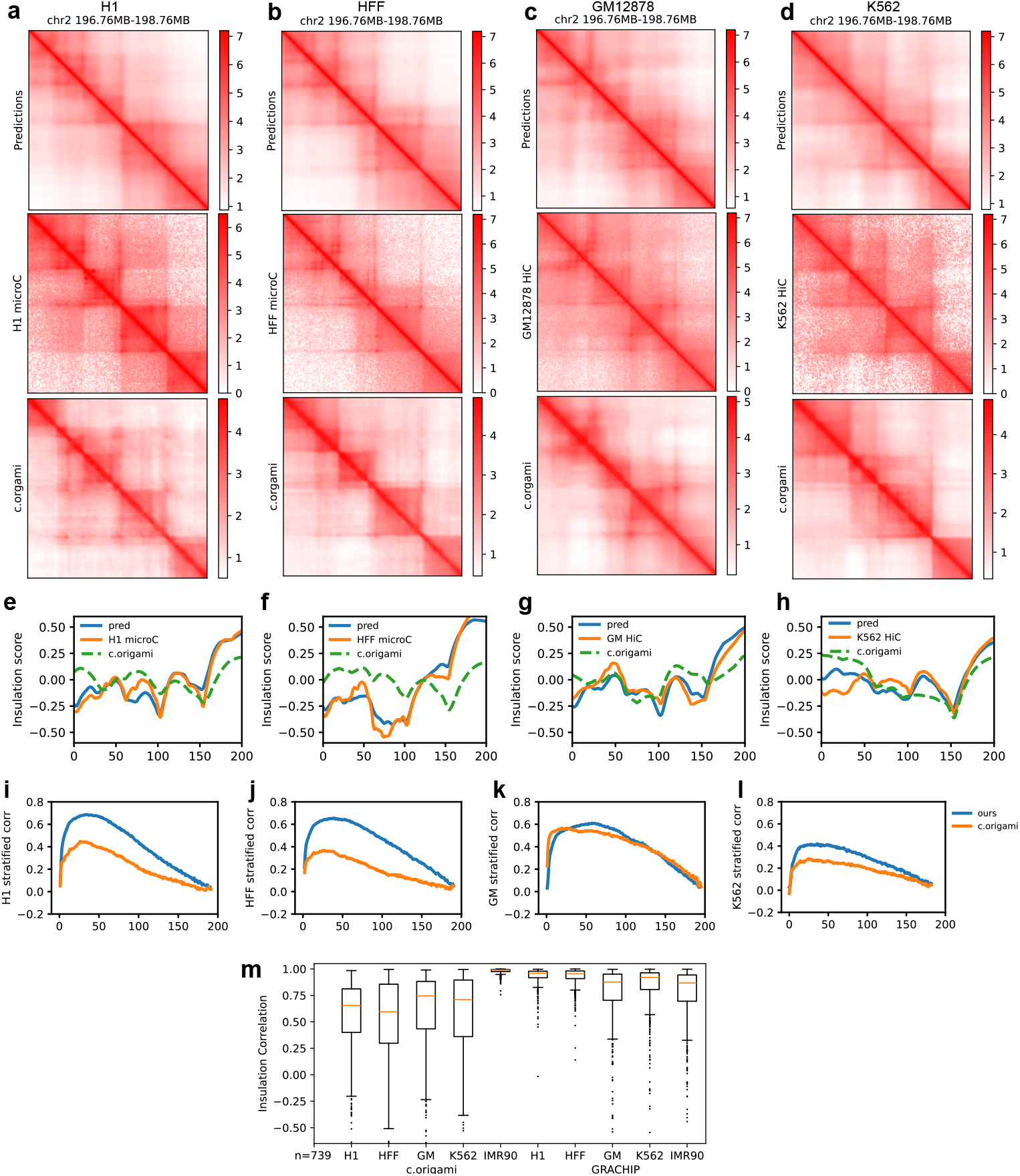
GRACHIP provides cell-type-specific interaction predictions. a-d, Examples of predicted genomic interaction (top), true genomic interaction (middle), c.origami provided interaction map (bottom) on H1, HFF, GM12878, and K562 cell line. e-h, insulation index of the example positions of predicted interaction maps, experimental interaction maps and c.origami provided interaction maps. i-l,distance stratified correlations between predictions and experimental maps, comparing GRACHIP to c.origami. m, Pearson correlations of predicted matrix insulation scores and true matrix insulation scores,comparing GRACHIP to c.origami on five different cell lines on chr2.

### Contributions of input chromatin contacts

Upon further investigation, it becomes evident that the proportion of chromatin contacts included in the input significantly influences GRACHIP’s performance, serving a dual function. Notably, the model achieves optimal performance when a higher proportion of input chromatin contacts (60%) is used, provided the prediction targets are derived from the same cell line as the input (Fig.4a). Conversely, a lower proportion (20%) is preferable when the prediction targets differ from the input cell line (Fig.4b-d). This observation suggests that an excess of input chromatin contacts, especially those incongruent with the prediction targets, can detract from the model’s focus on relevant input features, leading to an increase in false positive predictions and negatively impacting model generalizability. Remarkably, utilizing just 5% of input contacts enables GRACHIP to deliver cell-type-specific predictions with significantly higher accuracy than when employing 60% or more (Fig.4e-h), highlighting GRACHIP’s efficiency and its minimal reliance on supplementary information to surpass the performance of existing models.

**Fig. 4:**
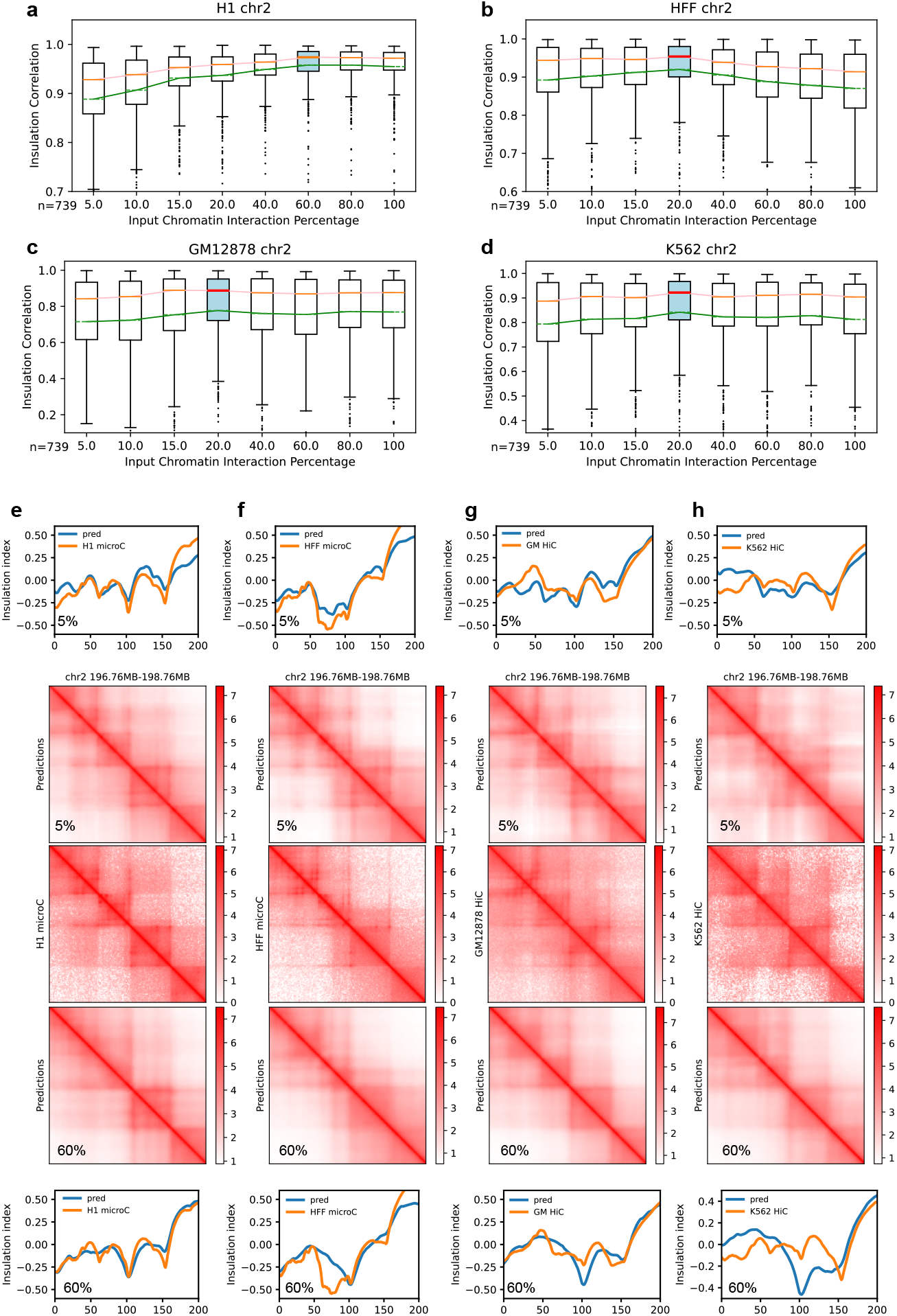
Input interactions play a dual role in the performance of the model. a-d, Pearson correlation of insulation scores grouped by different percentages of edges fed into the model. e-h, comparison of model performance with 5% and 60% input interactions.

Furthermore, GRACHIP’s performance on the H1 cell line does not uniformly improve with an increasing amount of chromatin contact information in the input (Fig.4a), potentially due to minor inconsistencies between the input contact dataset (H1 Hi-C) and the target dataset (H1 micro-C). This observation leads to the speculation that while chromatin contact information crucially enhances genomic interaction predictions—a novel approach adopted by GRACHIP—excessive information may detrimentally affect model performance and generalizability. Therefore, achieving an optimal balance between using chromatin contact information as a helpful hint versus as potential noise is paramount. In the context of our model, this balance is empirically established at a maximum of 20% input chromatin contact information, indicating a strategic threshold for maximizing GRACHIP’s predictive accuracy and applicability.

### Feature importance study of our model

We further conducted five sets of ablation experiments to assess the relative importance of each feature among the six genomic features used by our model, i.e., ATAC, CTCF, H3K4me, H3K4me3, H3K27ac, and H3K27me3, via ablation experiments. Specifically, we recorded the model performance by (1) disabling all but one feature (Fig.5a), (2) disabling one feature (Fig.5b), (3) disabling all but two features (Fig.5d), (4) disabling two features (Fig. 5e), and (5) disabling three features (Fig.5f). All the experiments are conducted by setting the ablated features to 0 at the inference stage, without re-training or fine-tuning the model. After experimenting with all the feature combinations, we aggregated the scores by the active features following the calculation of Shapley values [17] to quantify the contribution of each genomic feature to the high performance of GRACHIP (Fig.5g).

**Fig. 5:**
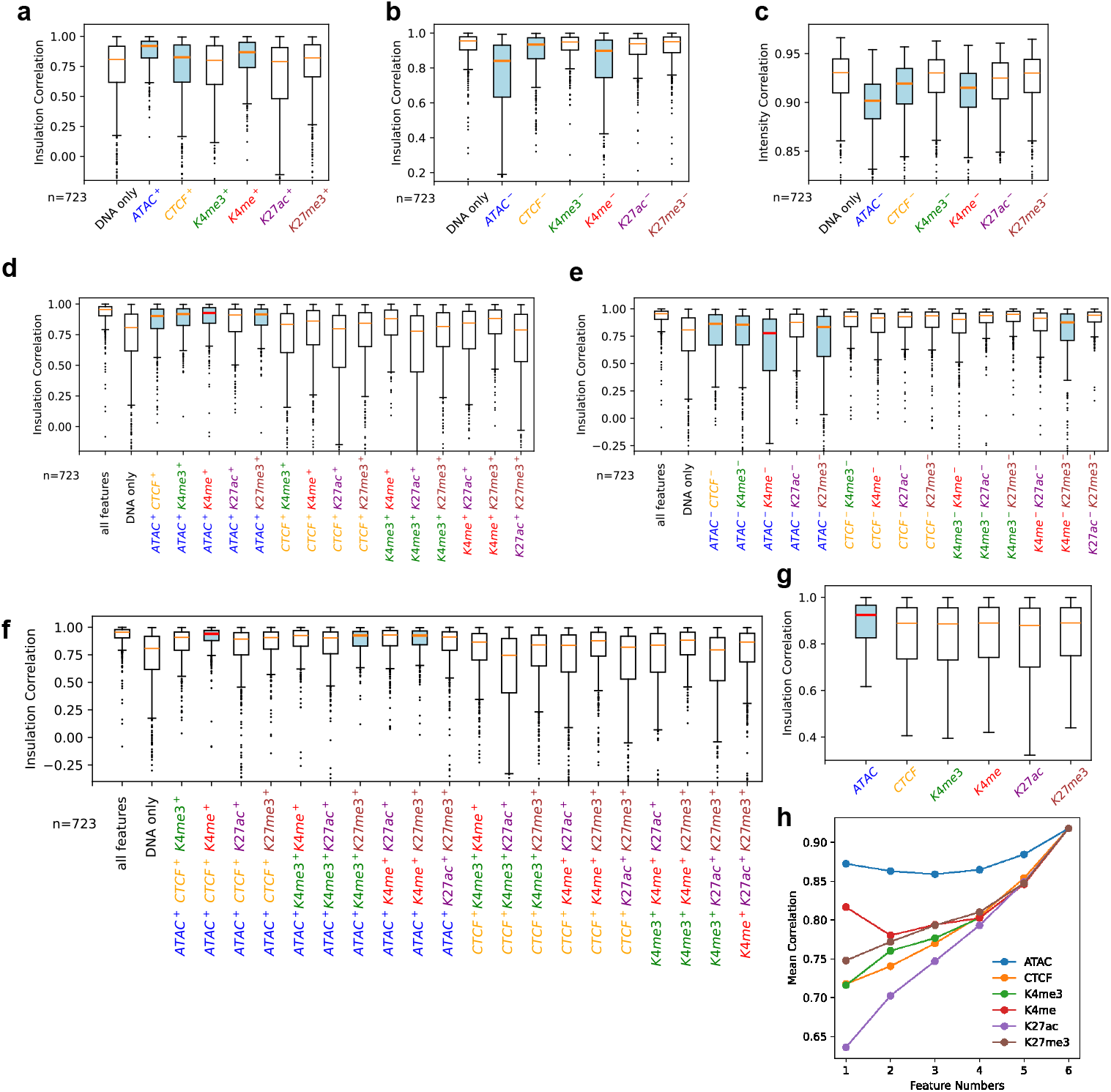
Feature ablation studies uncovered feature contribution. a, Pearson correlation of insulation score between predictions and experimental HFF microC. Examined the performance drop by disabling one feature. The disabled feature is marked as *Feature*^−^. b, Compare the performance of our model when disabling all the features but one. The remaining feature is marked as *Feature*^+^. d, Person correlation of insulation score when two features are remained. e, Person correlation of insulation score when two features are disable. f, Person correlation of insulation score when three features are remained. g. The average performance of predictions with specific features. h. Feature number stratified contributions evaluated by average all feature combinations’ insulation scores.

Among the six features, the ATAC-seq signal (accessible chromatin) stands out to be the most impactful feature of our model, as is consistent with previous research [8, 9]. H3K4 mono-methylation (K4me) is the second important feature, matching our expectation since K4me is a histone mark associated with promoters and primed enhancers [18], both of which play a crucial role in determining genomic interactions. While CTCF is ranked in the third place overall, its importance increases and surpasses that of H3K4me when more features are used in conjunction (Fig.5h). This observation, while also in alignment with existing literature, reveals another intriguing aspect of CTCF: on its own, CTCF binding may not provide sufficient information for predicting chromatin interactions. However, it has the potential to amplify the significance of other critical features or, conversely, be enhanced by them, thereby contributing more effectively to the predictive accuracy. To verify the accuracy of our feature rankings, we conducted additional experiments where the GRACHIP model was trained using three different feature sets: (1) ATAC and CTCF, (2) ATAC and K4me, and (3) ATAC, K4me, and CTCF (see Fig. S2). The results indicate that the combination of ATAC and K4me alone is sufficient to match the performance achieved with all six features. Moreover, configurations including K4me consistently outperform that with only ATAC and CTCF, underscoring K4me’s critical role in our model’s effectiveness.

Furthermore, larger feature groups (with three or more features) involving K27me3 exhibit decent contributions to the model’s performance (Fig.5f), which could be attributed to K27me3 being the sole marker representing super-repressive regions, making the information it provides irreplaceable. K27ac is shown to be the least significant feature (Fig.5g,h), likely resulting from the fact that regions marked by K27ac extensively overlap with those identified by other markers such as ATAC and K4me.

## Discussion

Enhancers can function cooperatively, with genetic variants within one enhancer influencing the activation of others, revealing a complex network of genomic interactions[11, 19]. Such cooperative functions can occur pairwise, between two promoters or between promoters and enhancers[12], as well as in multipartite arrangements among multiple cis-regulatory regions, commonly referred to as enhancer hubs or enhancer communities[11, 19, 20]. Inspired by these observations, our proposed model employs GNNs to capture the nuances of enhancer cross-talk, thereby improving its predictive accuracy for cell type-specific genomic interactions. Specifically, we utilize a dataset of known interactions derived from Hi-C experiments on the H1 cell line as a benchmark in our studies. While interactions identified in one cell line may not universally apply to others, they can still offer valuable insights into the model, as interaction regions tend to exhibit high correlation across different cell types.

Despite the benefits of incorporating known genomic interactions into our model, an excess of information specific to one cell type may introduce bias and negatively impact the model’s performance in other contexts. The model could erroneously rely on genomic interactions that are not present in the target cell lines, leading to false positive predictions. To address this, we conducted several experiments with varying proportions of input edges and empirically determined that using 20% of the edges from the H1 cell line optimizes performance. The methodology and outcomes of these experiments are detailed in the Results section.

The CNN architecture for GRACHIP has been carefully optimized for performance across five distinct cell lines. We opted for 2D CNNs over 1D CNNs because the latter tended to generate outputs with excessive noise, including numerous incorrect horizontal and vertical lines. Through extensive testing ranging from two to eight layers, we observed that deeper networks are more adept at capturing coarse-grained patterns, albeit at the expense of finer details. To strike a balance between capturing overall patterns and maintaining detail resolution, we settled on a configuration of six CNN layers. Additionally, incorporating max pooling layers between the CNN layers proved effective in reducing spurious horizontal and vertical lines in the predictions. Therefore, GRACHIP ultimately utilizes a configuration of six CNN layers interspersed with max pooling to optimize the quality of its outputs.

The pre-training methodology of GRACHIP distinguishes itself in two fundamental ways: through the novelty of its training paradigm and the strategic re-balancing of its training dataset. As delineated in the Training and Evaluation Strategies subsection within the Methods section, our approach introduces a graph-level objective for the recovery of chromatin interaction patterns. This objective is designed to foster model proficiency in reconstructing broad visual patterns evident in the actual data, in parallel to a microscopic focus on regression objectives for individual chromatin interactions. From a dataset perspective, we implemented a re-balancing strategy by downsampling instances that were overly consistent across two specific cell lines (H1 and HFF). This was intended to compel the model towards a finer recognition of inter-cell-line variances, notwithstanding identical DNA sequences, thereby enhancing its ability to generalize to novel scenarios. Despite a reduction in the volume of training data due to this subsampling, a significant uplift in model performance was observed, both for the cell line integral to the training process and for additional, untrained cell lines. This improvement underscores the efficacy of the applied training methodologies, with comprehensive results and analyses furnished in the Results section. Meanwhile, removing chromatin interaction intensity signals from the input markedly diminishes the model’s performance across all datasets, in stark contrast to scenarios where a minimal portion (5.0%) of ground-truth edges from the H1 cell line is incorporated as a hint. This outcome validates our hypothesis that even minimal ground-truth chromatin interaction intensity information can significantly enhance the accuracy and extend the generalizability of chromatin interaction prediction models, irrespective of the direct correlation between cell lines.

In summary, GRACHIP has proven its efficacy as a predictive model for chromatin interactions, validated through comprehensive assessments across various cell lines. The model’s robust performance on cell lines significantly divergent from its training data underscores its exceptional generalizability to other datasets or cell lines, without the need for further training or tuning. This characteristic establishes GRACHIP as a promising tool for efficiently testing new biological hypotheses, circumventing the need for expensive experiments to acquire chromatin conformation data.

## Methods

### Data processing

ATAC-seq, CUT and RUN, HiC, and MicroC datasets can be accessed through the ENCODE data portal (www.encodeproject.org) and the 4DN data portal (data.4dnucleome.org). Hi-C and MicroC data are processed following the 4DN Hi-C processing pipeline. Sequencing reads are mapped to the Hg38 human genome. CUT&RUN data are processed following 4DN CUT&RUN Processing Pipeline. ATAC-seq were processed following ENCODE ATAC-seq Processing Pipeline. Peak detection for ATAC-seq and ChIP-seq or CUT&RUN is performed with macs2[21]. The signals are then transformed to a log fold change format by macs2[21]. For HiC and MicroC data, matrices are normalized employing the ICE method[22] and subsequently converted to a natural log scale. Insulation scores are calculated after converting the contact maps back to the normal scale, adhering to the insulation score calculation method described in a prior study[23]. The entire genome is segmented into 10kb bins, and regions listed in the ENCODE blacklist[24] are excluded from the training set. This approach ensures that problematic regions known to introduce biases in genomic analyses are omitted, thereby improving the accuracy and reliability of the predictions. For predictive modeling, 200 consecutive bins are aggregated to form a 2MB chunk, which serves as the target unit for prediction.

### DNA sequences encoding

Given the established significance of DNA sequence information in identifying chromatin interactions, our model integrates DNA sequence encodings as a crucial component of its input. We specifically employ the pre-trained DNABert model to transform DNA sequences into 768-dimensional vectors, optimizing both computational efficiency and the depth of information extracted from the sequences, surpassing the capabilities of conventional one-hot DNA encodings. It’s noteworthy that in the implementation of GRACHIP, the DNABert model is utilized in its original form, with no adjustments made to its weights. However, there exists the potential to enhance model performance through the simultaneous optimization of DNABert weights during the pre-training phase. Future investigations may further explore the benefits of specifically training DNA encoding models in conjunction with chromatin interaction prediction tasks.

### Model structure

As depicted in Fig. 1, the proposed model consists of a 3-layer GCN module succeeded by a 12-layer 12-head Transformer encoder module and a six-layer CNN block. **Each GCN layer** takes as input the representation of each node in a graph (i.e., the aggregated DNA sequence information and genomic feature for each genomic region in our datasets) and node connectivity information (i.e., the interaction intensity information across genomic regions). Following Kipf and Welling [25], the genomic region representations are updated using the formula

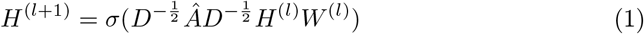

where *Â* = *A* + *I*_*N*_, *A* is the adjacency matrix indicating the interaction intensity between genomic regions in the input graph, *I*_*N*_ is the identity matrix of size *N* × *N*, 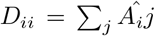 represents the degree of each node, *H*^(*l*)^ refers to the representation of each node as input to the current layer, and *W* ^(*l*)^ denotes the trainable weights at Layer *l. σ* denotes the sigmoid activation function. At the first layer of GCN, the input *H*^(0)^ = *W*^*feature*^*v*_*feature*_ + *v*_*DNA*_ + *PE* where *v*_*feature*_ and *v*_*DNA*_ are the genomic feature vector and DNABert encoding of each genomic region, respectively, *PE* represents the positional encoding, and *W*^*feature*^ is a trainable weight projecting the feature vector to the same dimension as the DNA vector (768). Positional encoding is calculated following BERT [15] using the following formula

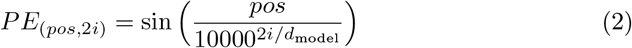

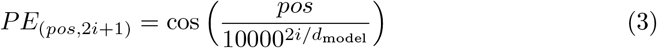

where *pos* represents the position of the genomic region in the sequence, 0 *≤ i ≤* (*d*_*model*_ − 1) represents the dimension, and *d*_*model*_ represents the dimensionality of the encodings. The update of genomic region encodings according to their neighbors’ encodings is illustrated in Fig. 1b. **The Transformer encoder block** takes the structure shown in Fig. 1c, where the *i* − *th* attention head on Layer *l* calculates self-attention following the formula

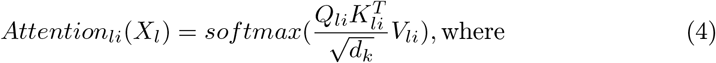

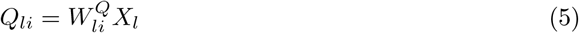

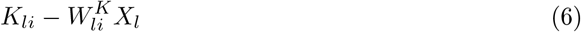

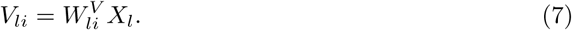

Here, 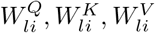 are trainable weights, *X*_*l*_ denotes the input representation to Layer *l*, and *d*_*k*_ is the dimensionality of hidden vectors *K*. Succeeding the calculation at each attention head, the outputs are aggregated for Layer *l* to become its output by

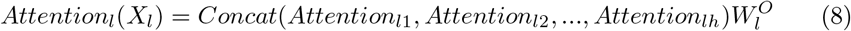

where *h* denotes the number of attention heads on Layer *l* (set to 12 in the proposed model) and *W*^*O*^ is a trainable weight. This output of intermediate Transformer layers (i.e., 0 *≤ l ≤* 10 in our model) is forwarded to the next Transformer layer as input after the GeLU activation is applied. No activation is applied to the output of the last Transformer layer. Similar to the GCN module, positional encoding is added to the input to the first layer of the Transformer by *X*_0_ = *H*^(3)^ + *PE* where *PE* denotes positional encoding and *H*^(3)^ is the output of the top layer of the GCN module.

**The CNN encoder block in our model** consists of six 2D convolutional layers designed to extract localized features from the hidden representations of genomic-region pairs. Each 2D convolutional layer employs *K* = *n*_output channel_ filters with a kernel size of 3*×* 3. At each position (*i, j*) of the input signal, each filter *k* spans over all the channels of the input, computing the convolution of the filter with the input signal segment of corresponding size across all input channels. This results in a feature map where the output at each position (*i, j*) for each filter *k* is a scalar *v*_*kij*_, computed as the sum of element-wise products between the filter weights and the input segment. These feature maps are then stacked across all filters to produce a composite output matrix *V*_*ij*_ = Concat(*v*_1*ij*_, *v*_2*ij*_, …, *v*_*kij*_) at each position (*i, j*). The convolution operation slides the filters across the entire input area, with a stride of 1 and padding set to 1 on all sides, producing an output matrix *V* of size *L × K*, where each feature vector at position (*i, j*) aggregates outputs from all filters.

In our proposed model, 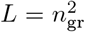 where *n*_gr_ = 200 denotes the number of genomic regions within each input instance. The input to the first CNN layer is of size *L×* (2 *×* 768), reflecting 1,536 channels by concatenating Transformer encodings of each pair of genomic regions. Subsequent CNN layers accept the outputs of their preceding layers as inputs, after applying the GeLU activation function. The number of output channels in the six CNN layers is progressively reduced: 768, 384, 192, 96, 48, and finally 1, gradually refining the dimensionality of the genomic-region-pair representations to a singular value.

The final output of the CNN encoder is reshaped to match the dimensions of the ground truth labels, which in our setting is 200 *×* 200.

In GRACHIP, the role of the GCN module primarily lies in encoding the reference genomic interaction information provided in the input, setting our model apart from other feature-only genomic interaction prediction models. Prior research in the field of Computer Science and Machine Learning has shown the efficacy of GNNs for graph-based representation learning and predictions, rationalizing our use of a GNN network in the model. Our preference for GCN over other GNN architectures is based on our preliminary experiments where GCN outperformed other GNN networks like Graph Attention Networks (GAT) [26] and GATv2 [27] on the genomic interaction prediction task.

### Training and evaluation strategies

Our model is initially pre-trained on the H1 and HFF cell lines, then assessed through zero-shot evaluation in both in-domain contexts (using reserved evaluation datasets from the H1 and HFF cell lines) and out-of-domain scenarios (utilizing evaluation datasets from the GM12878 and K562 cell lines). To ensure integrity, all input edges are derived exclusively from the H1 HiC dataset, thereby preventing any inadvertent disclosure of target-dataset labels to the model.

Pre-training of our model incorporates two distinct loss functions: an embedding loss to capture overarching patterns of the ground truth and a mean squared error (MSE) loss aimed at accurate prediction of edge intensity between pairs of genomic regions. An encoder-decoder model based on CNN architecture is preliminarily trained on a subset of the H1 HiC dataset (specifically Chromosome 1) to encode and reproduce ground truth data. This encoder of this model is subsequently employed to calculate the embedding loss by encoding both our model’s predictions and the ground-truth labels.

The composite loss function during training is defined as ℒ= ℒ_*MSE*_ +((*E//*100) − (*e//*100)) ℒ_*emb*_, where *E* and *e* denote the total number of training epochs and the current epoch, respectively, with ℒ_*MSE*_ and ℒ_*emb*_ representing the MSE and embedding loss components. This strategy prompts the model to initially focus on learning general interaction patterns between genomic regions, gradually shifting towards improving the accuracy of individual predictions as training progresses.

After pre-training, the model undergoes evaluation on datasets it has not previously encountered, without any additional training or adjustments. Detailed outcomes of these evaluations are thoroughly discussed in the Results section, showcasing the model’s performance and generalization capabilities.

## Data availability

The datasets used in this paper are listed in Supplementary Table S1.

## Code availability

The code for GRACHIP is at https://github.com/Ruoyun-W/GRACHIP.

## Author contributions statement

R.W., W.M., X.W., and S.V. conceived the experiments. R.W., W.M., A.M., and S.S. conducted the experiments. R.W., W.M., A.M., and S.S. contributed to the GitHub repo. R.W. and W.M. analyzed the results. W.M. and W.M. drafted the manuscript. All authors reviewed the manuscript.

## Additional information

**Table S1:** Datasets used in this paper.

| Cell Line | Experiment Type | Database | ID |
| --- | --- | --- | --- |
| H1 | microC | 4DN | 4DNFI9GMP2J8 |
| H1 | ATACseq | 4DN | 4DNFICPNO4M5 |
| H1 | CTCF CUT&RUN | 4DN | 4DNES1RQBHPK |
| H1 | K4me CUT&RUN | 4DN | 4DNESRFWR5SV |
| H1 | K4m3 CUT&RUN | 4DN | 4DNESPE6J9FU |
| H1 | K27ac CUT&RUN | 4DN | 4DNESIMWCLF8 |
| H1 | K27m3 CUT&RUN | 4DN | 4DNES8TY5P5P |
| HFF | microC | 4DN | 4DNFIOYPL9EC |
| HFF | ATACseq | 4DN | 4DNFIZ9191QU |
| HFF | CTCF CUT&RUN | 4DN | 4DNESJGIALEC |
| HFF | K4me CUT&RUN | 4DN | 4DNESR96HCCM |
| HFF | K4m3 CUT&RUN | 4DN | 4DNES8ZSCTFJ |
| HFF | K27ac CUT&RUN | 4DN | 4DNESV6NQ665 |
| HFF | K27m3 CUT&RUN | 4DN | 4DNESNNHF597 |
| GM12878 | HiC | 4DN | 4DNFIXP4QG5B |
| GM12878 | ATACseq | ENCODE | ENCFF603BJO |
| GM12878 | CTCF CUT&RUN | 4DN | 4DNES6GVE8XZ |
| GM12878 | K4me CUT&RUN | 4DN | 4DNESLYCNMZP |
| GM12878 | K4m3 CUT&RUN | 4DN | 4DNES9CEMVIB |
| GM12878 | K27ac CUT&RUN | 4DN | 4DNESOOUQAN7 |
| GM12878 | K27m3 CUT&RUN | 4DN | 4DNES7W1FL5I |
| K562 | HiC | 4DN | 4DNFI18UHVRO |
| K562 | ATACseq | ENCODE | ENCFF077FBI |
| K562 | CTCF CUT&RUN | 4DN | 4DNESMSEE62M |
| K562 | K4me CUT&RUN | 4DN | 4DNES6VYFIZW |
| K562 | K4m3 CUT&RUN | 4DN | 4DNESV634K6K |
| K562 | K27ac CUT&RUN | 4DN | 4DNFIEYAZI72 |
| K562 | K27m3 CUT&RUN | 4DN | 4DNES TTCK612 |
| IMR90 | HiC | 4DN | 4DNFIJTOIGOI |
| IMR90 | ATACseq | ENCODE | ENCFF282RNO |
| IMR90 | CTCF ChIPseq | ENCODE | ENCFF105FHL |
| IMR90 | K4me ChIPseq | ENCODE | ENCFF221MJG |
| IMR90 | K4m3 ChIPseq | ENCODE | ENCFF376ZIM |
| IMR90 | K27ac ChIPseq | ENCODE | ENCFF699OAR |
| IMR90 | K27m3 ChIPseq | ENCODE | ENCFF366BVS |

**Fig. S1:**
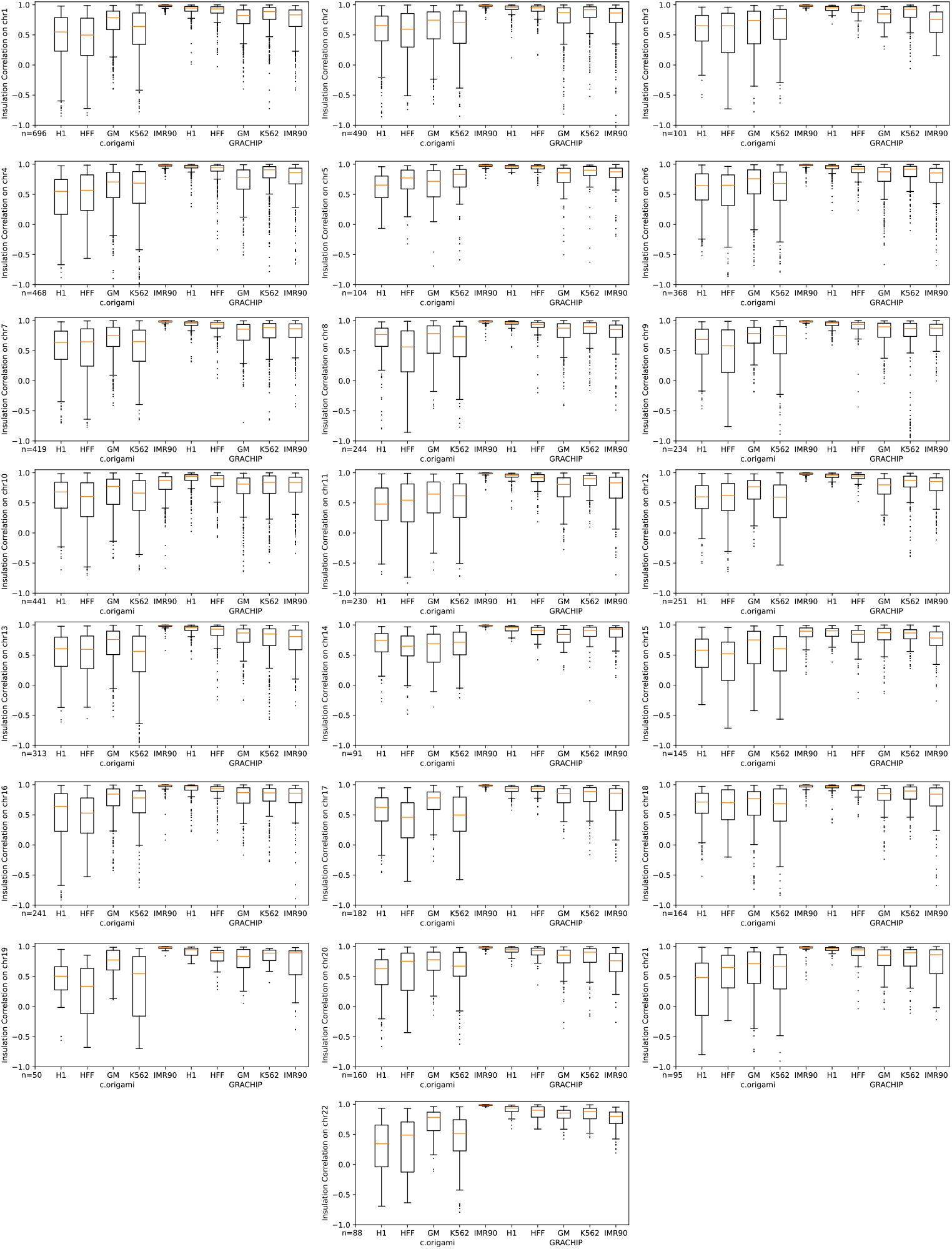
Whole-genome prediction performance of GRACHIP versus c.origami across five cell lines, quantified using Pearson correlation coefficients of insulation scores.

**Fig. S2:**
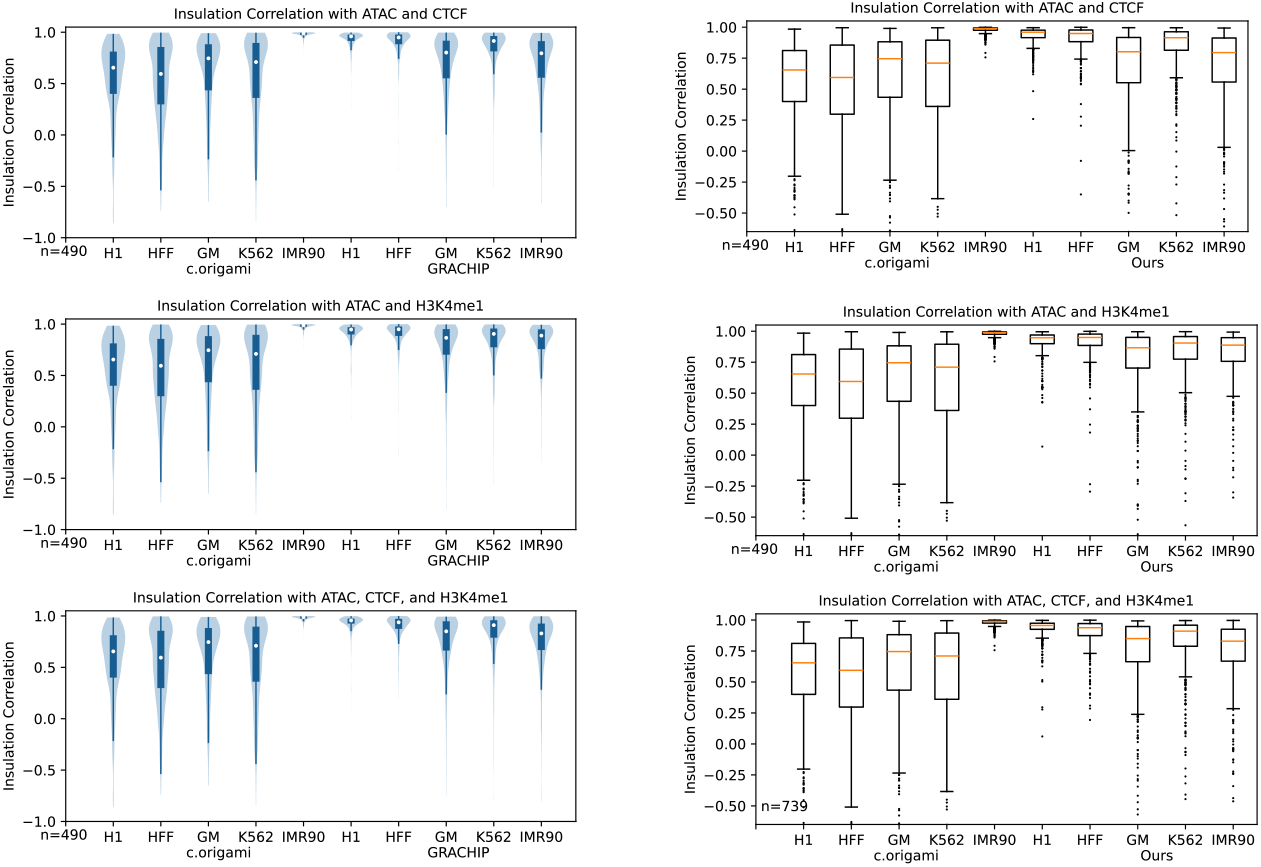
Comparison of insulation correlations between GRACHIP and c.origami using a subset of features, evaluated using Pearson correlation coefficients.

